# Precise modelling and correction of a spectrum of β-thalassemic mutations in human erythroid cells by base editors

**DOI:** 10.1101/2022.06.01.494256

**Authors:** Kirti Prasad, Nivedhitha Devaraju, Anila George, Nithin Sam Ravi, Gokulnath Mahalingam, Vignesh Rajendiran, Lokesh Panigrahi, Vigneshwaran Venkatesan, Kartik Lakhotiya, Yogapriya Moorthy, Aswin Anand Pai, Yukio Nakamura, Ryo Kurita, Poonkuzhali Balasubramanian, Saravanabhavan Thangavel, Shaji R Velayudhan, Srujan Marepally, Alok Srivastava, Kumarasamypet M Mohankumar

**Affiliations:** Centre for Stem Cell Research (a Unit of inStem, Bengaluru), Christian Medical College Campus, Bagayam, Vellore, 632002, Tamil Nadu, India; Manipal Academy of Higher Education, Manipal, 576104, Karnataka, India; Sree Chitra Tirunal Institute for Medical Sciences and Technology, Thiruvananthapuram, 695 011, Kerala, India; Department of Haematology, Christian Medical College & Hospital, Vellore -632 004, India; Molecular Cardiology Research Institute, Tufts Medical Center, 800 Washington Street Boston, MA, 02111; Cell Engineering Division, RIKEN BioResource Center, 3-1-1 Koyadai, Tsukuba, Ibaraki, 305-0074, Japan; Research and Development Department, Central Blood Institute Blood Service Headquarters, Japanese Red Cross Society, Japan

**Author notes:** Corresponding Author: Dr Mohankumar K. Murugesan, Centre for Stem Cell Research (a unit of inStem, Bengaluru), Christian Medical College Campus, Bagayam, Vellore-632002, Tamil Nadu, India.

**Keywords:** CRISPR/Cas9, Base editors, *HBB* gene deletion, CD34+ HSPCs, Human erythroid progenitor cells, β-thalassemia, HbE (CD26(G->A)), *HBB*-88(C->T), *HBB*-28(A->G), Initiation Codon(T->C), Codon 15(G->A), IVS1-5(G->A), IVS1-110(G->A), IVS2-849(A->G)

## Abstract

β-thalassemia and HbE result from mutations in the β-globin locus that impedes the production of functional β-hemoglobin and represents one of the most common genetic disorders worldwide. Recent advances in genome engineering have opened up new therapeutic opportunities to directly correct these pathogenic mutations using base editors that install transition mutations (A>G and C>T) in the target region with minimal generation of indels. Herein, for the first time, we demonstrate the usage of base editor in the correction of point mutations spanning multiple regions of the *HBB* gene, including promoter, intron and exon. To this end, we have engineered human erythroid cells harbouring the diverse *HBB* mutations, thus eliminating the requirement of patient CD34+ HSPCs with desired mutations for the primary screening by base editors. We further performed precise creation and correction of individual *HBB* point mutations in human erythroid cells using base editors, which were effectively corrected in the *HBB-engineered* erythroid model. Intriguingly, most bystander effects produced by the base editor at the target site were reported to exhibit normal hemoglobin variants. Overall, our study provides the proof-of-concept for the precise, efficient and scarless creation and correction of various pathogenic mutations at the coding and non-coding regions of *HBB* gene in human erythroid cells using base editors and establishes a novel therapeutic platform for the treatment of β-thalassemia/HbE patients. This study can be further explored in correcting the other monogenic disorders caused due to single base substitutions.

## Introduction

β-thalassemia represent one of the most common inherited monogenic disorders worldwide caused due to a spectrum of mutations in the β-globin gene, which results in a quantitative decrease in the production of functional β-globin chain or generation of structural hemoglobin (Hb) variants(1). The reduction or absence of β-globin chain synthesis caused due to mutations in the β-globin gene creates an imbalance in the α/β globin chain ratio leading to ineffective erythropoiesis(2). The degree of an imbalance created correlates with the severity of the phenotype, based on which β-thalassemia is categorised as β-thalassemia major (β^0^ / β^0^ - homozygous or as compound heterozygous with Hb structural variants), β-thalassemia intermedia (β^0^ / β^+^ - heterozygous) and β-thalassemia minor (β/ β^+/0^ - heterozygous)(3). At the molecular level, β-thalassemia consists of more than 350 disease-causing mutations comprising single nucleotide changes, deletions, and insertions(4). Most of the pathogenic point mutations reside in three major regions: the promoter and 5’ UTR, which cause defective β-gene transcription; the splice junction and polyadenylation sites, including the 3’ UTR, which interfere with normal mRNA processing; and the coding regions, which cause nonsense or frameshift mutation leading to defective mRNA translation(5). The only approved curative therapy available for β-thalassemia is an allogeneic stem cell transplant, which is often limited due to the lack of HLA matched donor availability(6). Ex-vivo genome editing offers an alternative therapeutic strategy wherein autologous transplantation of genome modified patient’s CD34+ HSPCs can be utilised for achieving a functional cure.

Gene editing strategies are focused on either the precise HDR or error-prone NHEJ mediated correction of the β-thalassemia mutations. Earlier studies used ZFNs, and TALENs followed by CRISPR/Cas-9 for HDR-mediated gene repair of several β-thalassemia mutations, like β-41/42(TCTT) deletion, *HBB*-28(A>G) and the highly prevalent HbE mutation (7–9). However, HDR-mediated gene repair is inefficient in primary CD34+ HSPCs, and the indel/HDR ratio is often very high, which might be detrimental in coding areas. Even with the high HDR efficiency achieved using small molecules, the long-term engraftment of the edited CD34+ HSPCs in mice is compromised which limits its therapeutic benefit (10–12). On the other hand, NHEJ based indel creation has been successfully attempted for the correction of intronic mutations such as IVS 1-110 and IVS 2-654, which creates aberrant splice sites (13,14). This approach is limited to intronic sites and cannot be used to correct mutations in the regions where it is crucial to preserve the sequence integrity, such as the coding sequences, promoter regions or splice sites. Nevertheless, the unintended effects of DSBs such as chromothripsis, large deletions, translocations, and p53 activation loom the prospects of Cas-9 mediated approaches (15,16). Therefore, the development of gene editing strategies for the correction of β-thalassemia mutations without induction of DSBs is crucial.

Base editors harness the advantage of the CRISPR–Cas9-based system by combining Cas9 nickase with either adenosine deaminase or cytosine deaminase, which converts A to G or C to T, respectively (17,18). The development of base editors will enable the efficient scarless single-nucleotide conversion at the desired target site that can avoid the generation of DSBs compared to other genome editing approaches, even in the non-dividing cells. This approach will overcome most of the limitations associated with the current genome editing platform based on HDR and NHEJ by creating the desired on-target mutation without needing a donor template and a double-strand DNA break. Base editors were found to be more preferential editing tools compared to Cas9 for the correction of disease-causing mutations in various monogenic disorders. The targeted disruption of the GATA1 binding site in the erythroid-specific enhancer region of *BCL11A* using CBE and recreation of various HPFH mutations in highly similar *HBG1* and *HBG2* promoters using ABE and CBE highlights the advantage of base editors in globin biology research (19–21). Most of the preclinical studies have reported minimal off-target effects at both DNA and RNA levels using base editors (22,23). An unavoidable consequence of base editing is the bystander effect, which might appear as a hurdle in correcting point mutations but could be circumvented to a great extent by careful design of gRNA and employment of Cas variants which recognise non-NGG PAMs (24,25).

Precise correction of β-thalassemia mutation requires the need for mutation-specific cellular models. The limited availability of patient cells for the preclinical studies due to disease complications emphasises the need to develop more relevant β-thalassemia disease models. Previous studies showed that erythroid cells differentiated from patient iPSC could serve as a disease model, but the generation, differentiation and characterisation of iPSC harbouring mutation are labour intensive and time-consuming (26,27). Prior findings successfully generated HbE/ β-thalassemia immortalised cell lines from the patient’s peripheral blood HSPCs (9). However, creating such cell lines for a spectrum of *HBB* mutations is less feasible. Base editing provides a direct single-step approach for mutagenesis and has been used efficiently to create various cellular and animal models of genetic diseases (28–30). Taking cues from the previous attempts, we generated a more relevant β-thalassemic disease model using base editors.

In the current study, we framed the strategy using base editors to precisely correct the most common pathogenic point mutations causing β-thalassemia. For the first time, we generated human erythroid cells harbouring a spectrum of β-thalassemia mutations. Using this model, we showed highly efficient and precise correction of multiple point mutations present in coding and non-coding regions of β-globin by base editors. The identified potential targets from the primary screening were further validated in the β-thalassemic cellular models harbouring specific β-thalassemia mutation. Overall, we demonstrated the higher efficiency of base editing for the modelling as well as correction of structural and functional β-globin variants in human erythroid cells. As the base editor mediated gene-editing approaches are entering the clinical trials, we anticipate that the approach we have demonstrated will have commendable therapeutic implications and benefit β-thalassemia patients in the near future.

## Results

### Engineering the human erythroid cells with a spectrum of β-thalassemia mutations

β-thalassemia is primarily due to the point mutations in the *HBB* locus, and efficient correction requires precise nucleotide conversions to reconstitute the functional hemoglobin production. We hypothesised that base editors could be used to directly correct β-thalassemia mutations. As a proof of principle, we evaluated the efficiency of base editors to correct the various key point mutations present in the promoter, Initiation Codon, intronic and exonic regions of the *HBB* gene that affect the different stages of β-globin production. The exact outcome of base editing is unpredictable and is often gRNA and locus-specific. In addition, a substantial difference in base editing efficiency that occurs within the editing window even in the same locus necessitates the preclinical validation of the designed gRNAs and base editor variants.

Therefore, first, we screened different variants of base editors for its potential in precise correction of β-thalassemia mutations with minimal generation of indels and bystander edits. For the primary screening, we generated human erythroid cells with a spectrum of β-thalassemia mutations **(Figure 1a)**, as it is often challenging to obtain a sufficient number of patient CD34+ HSPCs with desired point mutation. We incorporated the most prevalent Indian and global point mutations in a *HBB* gene fragment that could be targeted using base editors **(Figure 1b)** and integrated into the human erythroid cells by lentiviral transduction. Before incorporating the *HBB* gene fragment harbouring various β-thalassemia mutations, we deleted the endogenous *HBB* gene, including the promoter, from the HUDEP-2 cell line to avoid the potential bias in editing outcomes while validating the efficiency of guide RNAs. The gRNAs targeting 5’ and 3’ end of the *HBB* gene were first validated individually **(Supplementary figure 1a)** and then nucleofected simultaneously as Cas9 RNP complex into HUDEP-2 cells to delete the HBB locus. Next, we confirmed the deletion by PCR **(Figure 1c and Supplementary figure 1b)** and quantified the deletion by qRT-PCR **(Figure 1d).** The pool of *HBB* deleted homozygous clones from the single-cell sorting (5 clones) was expanded for further experiments (referred to as *HBB* null cells) **(Supplementary figure 1c)**. We performed a multiplex ligation-dependent probe amplification (MLPA) assay in the pool of cells, a commonly used method for analysing the copy number variations in β-thalassemia patients, which further confirmed the *HBB* deletion with no signals detected by the probes targeting the *HBB* locus **(Figure 1e).** As reported previously, we also noticed a compensatory increase in the number of fetal hemoglobin positive cells after *HBB* deletion (Data not shown) (31).

**Figure 1:**
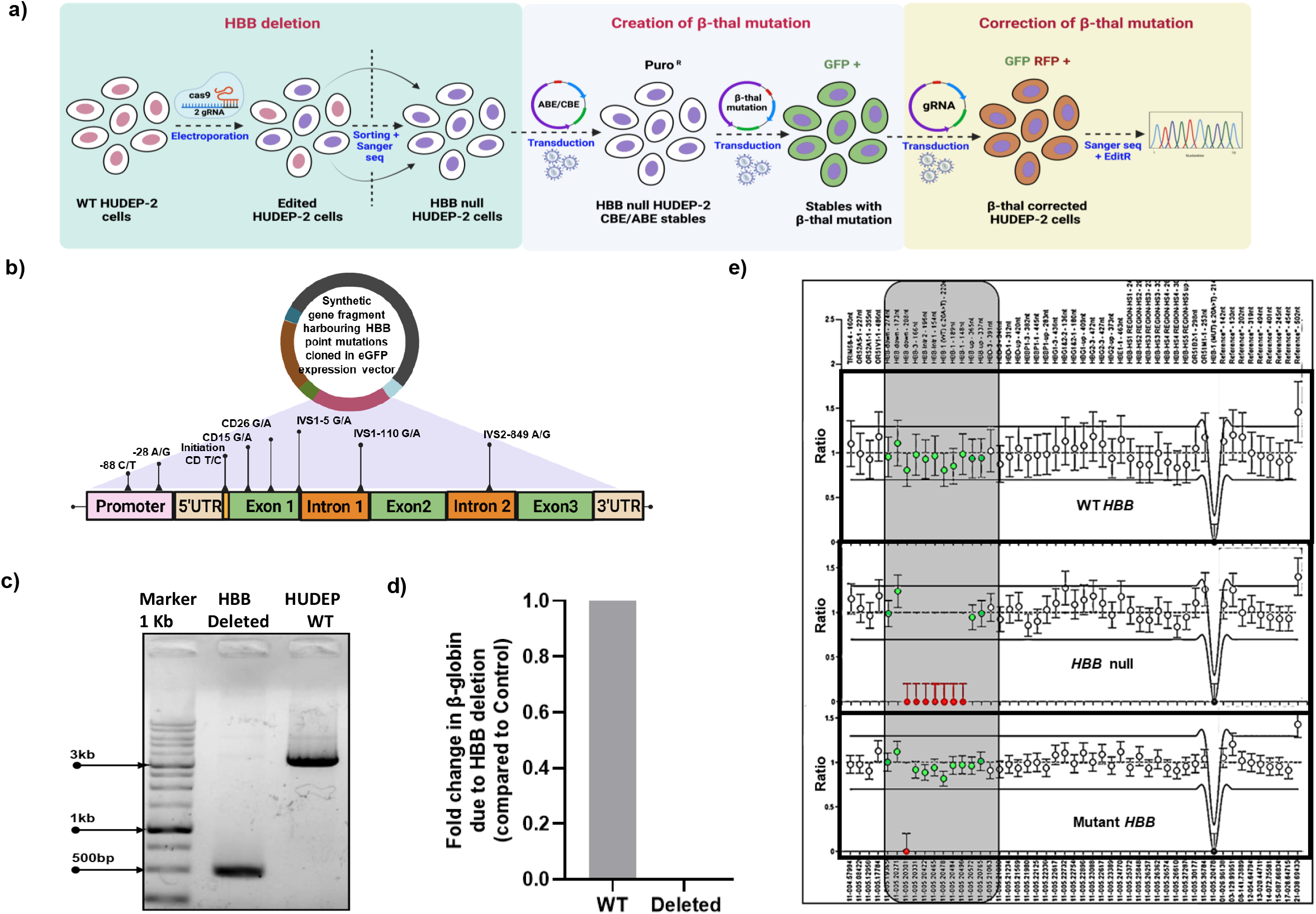
Generation of *HBB* null HUDEP-2 cells carrying spectrum of β-thalassemia point mutations: **a)** Schematic representation of creating engineered HUDEP-2 cells expressing base editor to validate gRNA for the correction of β-thalassemia point mutations. The engineered HUDEP-2 cells are devoid of the native *HBB* gene while harbouring an *HBB* gene fragment with desired β-thalassemia point mutations and expressing base editors. **b)** Layout of *HBB* gene fragment harbouring β-thalassemia point mutations targeted in this study are represented. **c)** Amplification of *HBB* gene in β-globin deleted HUDEP-2 cells and control; bands at 500bp and 3330bp represent the deleted and control samples, respectively. **d)** *HBB* gene deletion quantified in the genomic DNA using qRT-PCR in *HBB* deleted HUDEP-2 cells. **e)** Analysis of *HBB* region in HUDEP-2 cells before deletion and after deletion, insertion of mutated *HBB* gene fragment in deleted cell line using MLPA.

*HBB* null cells expressing ABE (ABE 7.10 and ABE 8e) or CBE were generated to facilitate the base editing for further experiments. Base editing potential of these stable cell lines was confirmed using a previously reported gRNA targeting the *BCL11A* binding site in the *HBG* promoter **(Supplementary figure 1d)**. These cells were then transduced with a lentiviral vector encoding the *HBB* gene fragment harbouring various β-thalassemia mutations and GFP reporter at 30% transduction efficiency to ensure single copy integration. MLPA analysis confirmed the successful integration of the gene fragment in the *HBB* null cells expressing GFP **(Figure 1 e**). Thus, we generated different base editor variants expressing HUDEP-2 cells with a spectrum of β-thalassemia mutations. We anticipate that the generated unique β-thalassemia cellular model would provide an efficient and robust platform for screening base editor variants in the correction of various β-thalassemia mutations.

### Primary screening of base editor variants for the correction of various β-thalassemia mutations in the *HBB* promoter, exons and introns

Utilising the engineered HUDEP-2 cells harbouring various β-thalassemic mutations, we evaluated the efficacy of base editor variants, on-target gRNA efficiency and bystander effect in the correction of selected pathogenic point mutations that affect *HBB* expression at several stages, involving transcription, mRNA processing and translation, or result in the production of structural variants. Target base substitution of adenine to guanine was investigated using two different variants of adenosine base editor (ABE 7.10 and ABE 8e), and cytosine to thymine conversion was evaluated through cytosine base editor. Depending upon the availability of PAM and the target nucleotide position within the editing window, we designed and validated unique gRNAs targeting specific *HBB* point mutations. All gRNAs were shown to have>90% transduction efficiency in engineered HUDEP-2 cells **(Supplementary figure 2b)**. *HBB* promoter comprises sequences from 100 bp upstream to the site of the initiation of transcription, including functionally important CACCC, CCAAT and ATAA boxes which are crucial for the binding of transcriptional factors that promote the expression of β-globin (1). Any point mutations that occur in promoter regions effectively reduce the production of *HBB* mRNA transcripts leading to β^+^phenotype (5). Therefore, we sought to correct the two prevalent promoter mutations, −88 (C>T) and −28(A>G) **(Supplementary figure 2a)**. We employed both ABE variants for the reversal of −88(C>T) mutation; ABE 8e exhibited 80% conversion at the −88 position with minimal bystander effects at −98 site **(Figure 2a)**. On the other hand, ABE 7.10 produced less than 20% on-target editing without any undesired edits, which might be attributed to the low enzymatic fidelity of ABE 7.10 compared to ABE 8e. Further, we employed CBE for the base correction at *HBB*-28(A>G) promoter mutation, which displayed an equivalent conversion efficiency at both on-target and bystander sites **(Figure 2a)**. Although the undesired *HBB* −25(G>A) conversion has not been reported to cause the β-thalassemia phenotype, another variant, *HBB* −25(G>C), has shown to be pathogenic, necessitating a better understanding of the *HBB* −25(G>A) conversion. These results indicate that careful selection of base editor variants and gRNAs may effectively correct the various *HBB* promoter mutations with the highest precision and efficiency.

**Figure 2:**
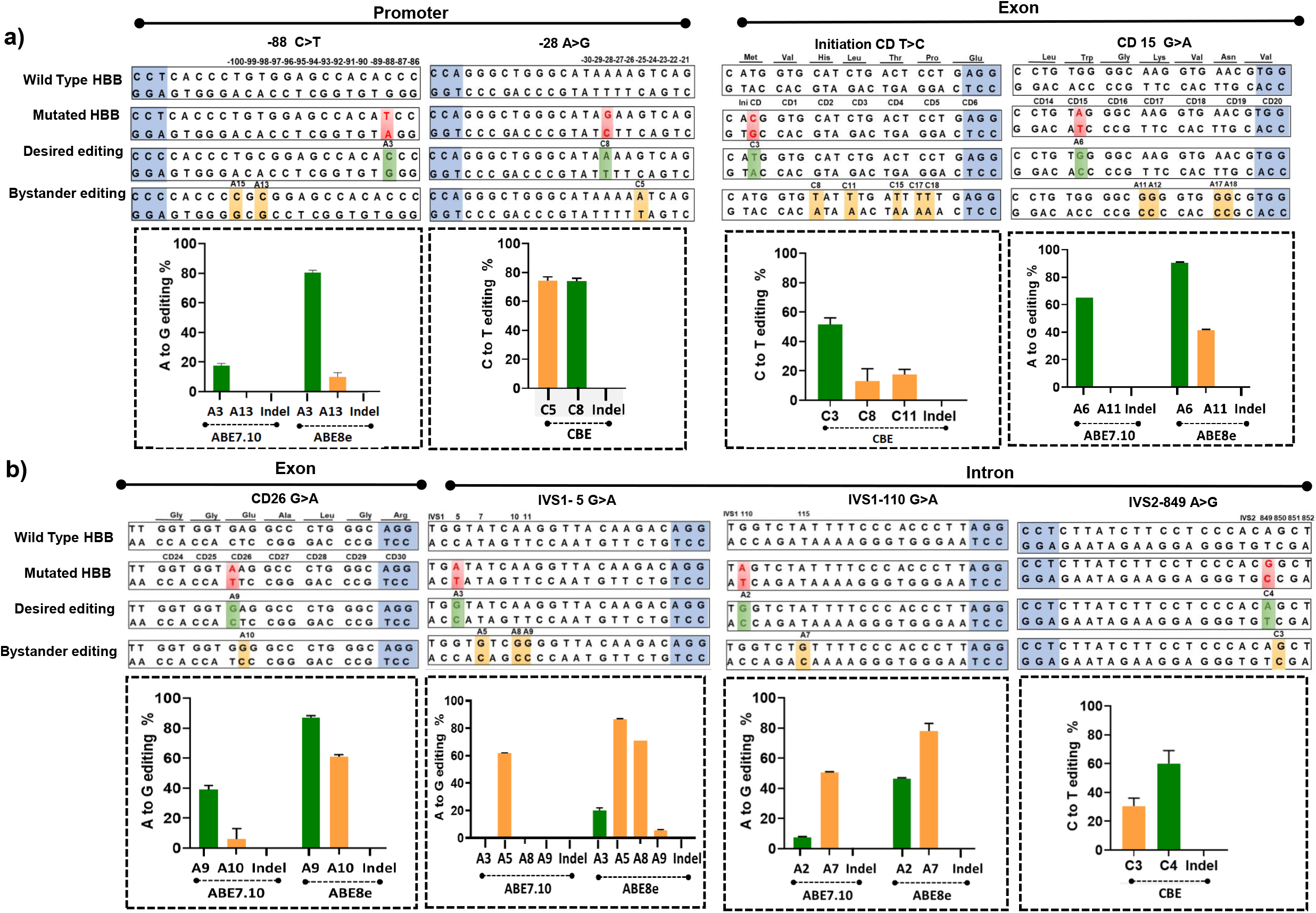
Correction of various β-thalassemia point mutations in the engineered HUDEP-2 cells by base editors. **a-b)** A graphical representation of the wild type and mutated *HBB* DNA sequence targeted by the respective mutations gRNA, mutated nucleotide (red), desired editing (green) and undesired bystander editing (yellow) are presented; highlights in blue represent the PAM sequence. The point mutations were classified into three categories based on their position in the *HBB* gene;1) Promoter region, 2) Exonic region, and 3) Intronic region. The desired point mutation in the engineered *HBB* locus was corrected using different variants of base editors. The substitution efficiency for eight different *HBB* point mutations is represented, where the green bar represents the intended substitution while the orange bar represents the unintended bystander editing. Data are expressed as mean ± SEM from two experimental replicates.

Next, we aimed to reverse the β^0^ phenotype caused due to mutations in the transcription start Codon of *HBB*. At least 8 point mutations originate from base conversion at any of the three nucleotides in the start Codon that affect the *HBB* mRNA translation(1). This study investigated the correction efficiency of the common Initiation Codon(ATG> ACG) mutation using CBE. We achieved 51% correction at the desired target site with a lower percentage of unwanted deamination in the *HBB* Codons 2 and 3 **(Figure 2a)**. Interestingly, previous clinical reports indicate the existence of non-pathogenic mutations coinherited with Initiation Codon(ATG> ACG). As a result, while editing Initiation Codon(ATG> ACG), we carefully investigated all conceivable bystander changes. We observed that the minimal bystander edits at Codon 2(CAT> TAT) created Hb Fukuoka - a normal variant in heterozygous form, whereas at Codon 3(CTG>TTG) causes a silent mutation, respectively **(Supplementary figure 2c)** (32).

Another important class of severe β-thalassemia mutations are the nonsense mutations within the *HBB* coding regions. We utilised two different variants of ABE to correct the frequent Asian/Indian mutation at *HBB* CD15(G>A), which creates a premature stop Codon and leads to mRNA degradation **(Supplementary figure 2a)**. To our surprise, we observed 65% and 90% editing to correct *HBB* CD15G>A mutation with ABE 7.10 and ABE 8e, respectively **(Figure 2a)**. However, most bystander edits had shown to result in a normal phenotype in heterozygous form. The most prominent bystander effect observed with ABE8e at CD17 changing the wild type (AAG, Lys) allele to Hb-Nagasaki (GAG, Glu) is a naturally occurring haemoglobin variant, and individuals with heterozygote genotype are shown to have normal haematological parameters **(Supplementary figure 2d)**. Although the on-target editing was marginally lower with ABE 7.10 at this locus, the editing at the non-target nucleotides was absent compared to modifications created by ABE 8e. These results emphasise the need for preclinical validation and careful selection of gRNA and base editor variants to correct pathogenic point mutations with minimised unwanted edits.

Furthermore, alterations that affect the processing of *HBB* mRNA transcripts result in severe/mild β-thalassemia phenotype, which is caused mainly due to the base substitution at the invariant dinucleotides (GT or AG) at the donor/acceptor splice sites and consensus flanking regions surrounding exon-intron junction (1). *HBB* IVS1-5(G>A) mutation that affects the consensus sequence, thereby decreasing the efficiency of conventional splicing **(Supplementary figure 2a)** (1). Here, we employed both ABE 7.10 and ABE 8e variants to correct the *HBB* IVS1-5 (G>A) mutation. Unlike the other two validated target regions, we identified that the bystander edits (at *HBB* IVS 1-7 and 1-10 positions) generated were more prominent (~4 fold) compared to on-target conversion at the IVS 1-5 site utilising ABE 8e while ABE 7.10 failed to correct the required base and only created bystander mutations in the *HBB* IVS 1-7 site **(Figure 2b)**. Noticeably, the bystander edit creates a point mutation outside the desired sequence within the intron, which might be dispensable for normal splicing.

We then aimed to address a cryptic splice site mutation, IVS1-110 G>A, predominantly in the Mediterranean, Middle East and Cyprus regions (33). This mutation is known to disrupt normal β-globin splicing by generating a de novo splice acceptor site in *HBB* intron-1 that leads to the production of an aberrant mRNA retaining 19 nucleotides from the intron prior to the start of exon 2, resulting in an in-frame premature stop codon **(Supplementary figure 2a)** leading to severe β-thalassemia with less than 10% of normally spliced mRNA transcripts (14). Previous attempts to rectify this mutation using CRISPR Cas 9 based non-specific disruption of the cryptic splice site have restored the normal splicing machinery. However, in this study, we employed specific corrections using ABE. At the IVSI-110(G>A) site, ABE 7.10 delivered lower editing due to the location of the target base outside the editing window, but the high-fidelity ABE 8e variation generated a 50% conversion **(Figure 2b)**. Although utilising ABE 8e improved editing efficiency, base conversion at IVSI-115(A>G) was unavoidable because it was within the editing window from the PAM site. Based on the earlier results of Cas9 induced non-specific disruption at this region, we believe that the created intronic mutation at *HBB* IVSI-115(A>G) may not be functionally relevant, but more in-depth studies are required to establish the functional implications of the unintentional edit. Next, we attempted to correct the common intron-exon splice junction mutation, *HBB* IVS 2-849(A>G), prevalent in African Americans. The effort to correct the *HBB* IVS 2-849 (A>G) mutation using CBE resulted in 60% on-target conversion along with bystander editing of 30% at *HBB* IVS 2-850 (G>A) at the splice junction **(Figure 2b)**. Even though half of the modifications are with bystander effects, we anticipate that the remaining G to A conversion at *HBB* IVS 2-849 would be sufficient to restore functional β-globin production.

Finally, we attempted to correct the most frequent pathological structural variant, the HbE CD26(G>A) mutation, highly prevalent in Asia, resulting in a very severe disease phenotype inherited as compound heterozygous HbE/β-thalassemia (34). HbE/β-thalassemia accounts for one-half of all severe β-thalassemia among all β-thalassemic mutations (35). Correction of the HbE mutation alone is enough to reduce the severity of compound heterozygote HbE/β-thalassemia symptoms and clinical manifestations. We attempted to reverse the HbE, CD26(G> A) point mutation using ABE as it is caused by a transition point mutation. We achieved the 39% and 87% on-target conversion of HbE CD26(G>A) mutation using ABE 7.10 and ABE 8e, respectively **(Figure 2b)**. Although the bystander effect was observed for both the variants of ABE, the on-target editing was 8-fold higher while using ABE 7.10 and ~2-fold higher when edited with ABE 8e. Nevertheless, the bystander edit resulted in the creation of Hb-Aubenas (CD26 AAG > GGG (Lys>Gly)), which is shown to be a naturally occurring hemoglobin variant with normal hematological and clinical features in heterozygous conditions **(Supplementary figure 2e)** (32). The new variant generated due to bystander edit at CD26 AAG > AGG (Lys>Arg) has to be examined further.

Although we observed several unintended bystander edits while utilising base editors for gene correction, we did not identify any insertions or deletions (InDels) in sanger sequencing results which are a common by-product of Cas 9 mediated editing. We found that most bystander edits generated during gene correction with the chosen gRNA and base editor variant resulted in a normal phenotype or were not implicated in *HBB* functional expression except in the intronic region. Collectively, our data demonstrate a base editor-mediated restoration approach for multiple β-thalassemia point mutations located in the promoter, exonic, and intronic regions of the *HBB* gene using engineered HUDEP-2 cells harbouring various β-thalassemia mutations.

### Creation of a human erythroid cellular model of β-thalassemia /HbE using a base editor and Cas9

Though we demonstrated a successful primary screening of base editor variants for the correction of various β-thalassemia mutations in the engineered *HBB* constructs, the functional assessment of its effects on restoration of β-chain synthesis needs to be validated in the native locus, which requires precise modelling of the various β-thalassemia mutations in human erythroid cells. As base editors are shown to be efficient and accurate in scarless incorporation of point mutations, we created two different severe β^0^-thalassemia mutations at the coding (Initiation Codon(T>C)) and intronic (IVS2-849(A>G)) region of *HBB* locus in HUDEP-2 cells.

We delivered ABE 8e mRNA and the respective gRNA by nucleofection to create Initiation Codon (T>C) and IVS2-849(A>G) mutations in HUDEP-2 cells **(Figure 3a)**. The overall editing efficiency for the generation of *HBB* Initiation Codon(T>C) mutation was high but was accompanied by undesired bystander edits **(Figure 3c)**. Nevertheless, single-cell sorting of the population yielded clones with homozygous intended edits, heterozygous edits and clones with unintended edits.

**Figure 3:**
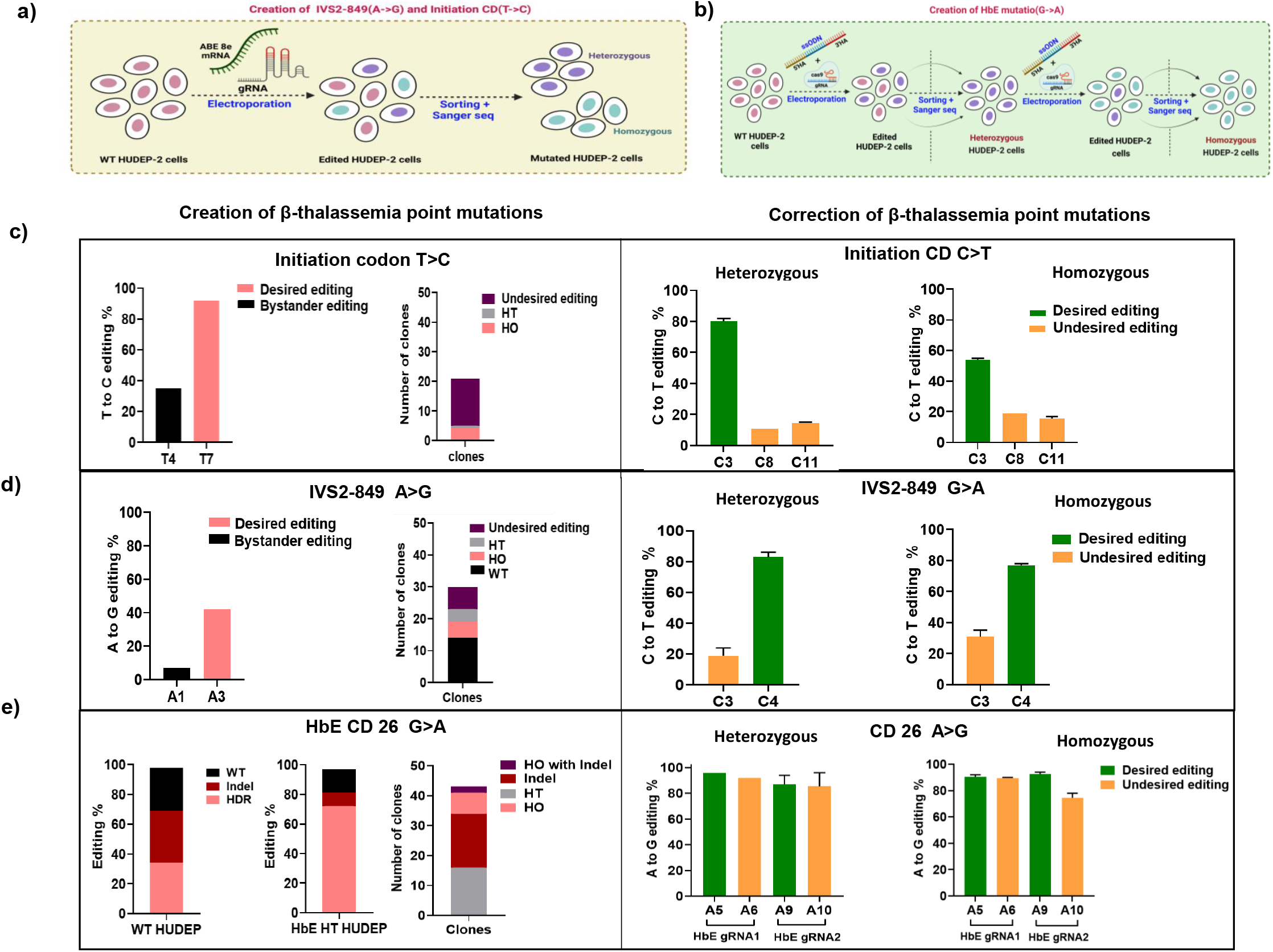
Creation and correction of Initiation Codon(T->C), IVS2-849 (A->G) and HbE (G->A) pathogenic mutations in HUDEP-2 cells using base editor and Cas9: **a & b)** Graphical representation for the creation of Initiation Codon(T->C), IVS2-849 (A->G) using ABE (a) and HbE using HDR (b). **c-e) Creation of β-thalassemia/HbE point mutations in HUDEP-2 cells:** Once the desired mutations were confirmed using Sanger sequencing (images in the left), single-cell sorting was carried out to obtain homozygous and heterozygous mutant clones. The clones were then validated by Sanger sequencing; the images on the right represent the number of clonal variants obtained. **c-e) Correction of β-thalassemia/HbE point mutations in HUDEP-2 cells:** The homozygous and heterozygous mutant clones with the desired mutation were pooled and transduced to express the base editors. The images represent the desired correction and undesired bystander edits in heterozygous (in the left image) and homozygous disease genotypes (in the right image). Data are expressed as mean ± SEM from two experimental replicates.

During the creation of *HBB* IVS 2-849(A>G) mutation, we observed a relatively low level of on-target desired nucleotide conversion with minimal bystander effects **(Figure 3d)**. Even though we obtained a mosaic combination of edited erythroid cells, single-cell sorting yielded a significant number of clones with homozygous and heterozygous variants for *HBB* IVS 2-849 (A>G) point mutation.

In addition to creating two severe β-thalassemic mutations, we also attempted to create a functional model for the most common pathological structural hemoglobin variant, HbE CD26 (G>A) mutation in human erythroid cells. We employed an HDR based approach using Cas-9 and ssODN to insert HbE CD26(G>A) mutation as the lack of a suitable editing window and PAM availability limits the use of the base editor. However, the lower efficiency of HDR **(Figure 3e)** mediated creation of the HbE CD26(G>A) mutation makes it difficult to acquire homozygous mutation even after single-cell sorting. Therefore, unlike creating other *HBB* mutations using a base editor in a single step, we adopted a two-step approach to creating HbE CD26(G>A) mutation with HDR **(Figure 3b)**. To accurately simulate the HbE CD26(G>A) mutation, in the first step, heterozygous clones were generated, followed by editing in these heterozygous populations to achieve homozygous clones with different gRNA **(Figure 3e)**. The edited cells were again sorted into individual cells to isolate clones with homozygous and heterozygous for HbE CD 26(G>A) mutation. These results suggest that base editors are highly efficient in the precise creation of a cellular disease model for β-thalassemia/HbE mutation in human erythroid cells. These β-thalassemic cellular models will be used to efficiently correct the specific *HBB* point mutation and validate for the functional restoration of adult hemoglobin tetramer (α2β2) by HPLC.

### Correction of mutations in human β-thalassemic cellular models using base editors

Next, we evaluated the efficiency of potential gRNAs and base editor variant that we have identified from preliminary screening on the correction of *HBB* mutations in the β-thalassemic cellular models. CBE-expressing HUDEP-2 stable cells were generated from *HBB* Initiation Codon (ATG>ACG) and *HBB* IVS 2-849(A>G) mutations by lentiviral transduction. The efficiency of gene correction in the β-thalassemic cellular model transduced with appropriate gRNA was evaluated by Sanger sequencing. Gene correction of the Initiation Codon (ATG>ACG) mutation using CBE resulted in 80% and 54% editing at the target base in clones of the heterozygous and homozygous cellular model with less than 20% bystander edits at two adjacent codons (Codon 2 and 3) with no detectable insertions or deletions which were consistent with the findings obtained in the preliminary screening using engineered *HBB* construct **(Figure 3c)**.

Similarly, we performed base editor-mediated gene correction of *HBB* IVS 2-849(A>G) mutation in a generated cellular model. We observed cytosine deamination at both sites as the desired base ‘C’ at IVS 2-849 to be corrected, followed by another ‘C’ at IVS 2-850 position, and both the bases were within the efficient editing window. Interestingly, we found a higher intended on-target conversion at *HBB* IVS 2-849(A>G) in the native locus and the designed *HBB* construct with a modest bystander impact **(Figure 3d)**. Given that the bystander effect accounted for roughly 30% of total editing that might influence the consensus sequence at the splice junction, the remaining cells, around 50% only with the desired editing, will be sufficient to contribute to normal splicing restore the functional β-globin chain production.

In addition, we also attempted to reverse the HbE genotype in the HUDEP-2 cells carrying the *HBB* Codon 26(A>G) mutation and expressing ABE. For the reversal of the missense mutation at *HBB* Codon 26( A>G), we used ABE variants; however, the efficient editing window also permitted the conversion of ‘A’ at the second position in *HBB* Codon 26. Bystander editing was unavoidable due to desirable and undesired bases adjacent to one other, resulting in equal modification at both sites **(Figure 3e)**. Surprisingly in HbE mutation, the bystander edits, which results in Hb Aubenas *HBB* Codon 26 (Glu>Gly) phenotype, gained predominance when tested in the native locus, which was reported to exhibit normal phenotype. However, we did not observe any undesired insertions or deletions at the target site after the gene-editing was analysed by sanger sequencing.

In conclusion, we achieved comparable on-target editing efficiency and bystander effects in both the engineered *HBB* construct and β-thalassemic cellular model, which underlines the merit of using the human erythroid cellular model harbouring spectrum of β-thalassemia mutations for the primary validation of base editor variants. Utilising a base editor in the β-thalassemic cellular model that mimics the genotype with an actual patient sample, we were able to successfully demonstrate the reversal of the β-thalassemia/HbE mutation and the restoration of β-hemoglobin will be validated by HPLC.

## Discussion

Genome editing approaches has become an attractive strategy for the treatment of β-thalassemia as even partial correction of disease-causing mutation is sufficient to restore the functional hemoglobin production and ameliorate the severity of the disease. Most of the current studies rely on Cas9 dependent HDR mediated direct correction of various β-thalassemia mutations. The major drawback of the CRISPR-Cas9 HDR system for disease correction is its relative inefficiency in quiescent cells and the creation of large unintended deletions and chromosomal-level alterations due to the DSBs (16). Moreover, insertions and deletions at the coding region of the β-globin gene might lead to potentially severe β-thalassemia major phenotype.

Previous studies have demonstrated the use of base editors to create or correct specific mutations in different genomic loci in various genetic disorders (28,36–38). Most studies in β hemoglobinopathies have demonstrated the therapeutic up-regulation of fetal globin expression by introducing HPFH-like mutations or by disruption of fetal globin repressors using base editors to compensate for the deficient β-globin (23). Despite this, only a few attempts have been performed to address the correction of β-thalassemia point mutations using a base editing strategy. The most common *HBB* −28(A>G) promoter mutation that decreases the production of β-globin has been extensively studied for correction using different base editor variants in various cell types (19,39). Further, electroporation of cytosine base editor and gRNA as RNP complex corrected the *HBB* −28(A>G) mutation with high efficiency and minimal bystander edits (19).

In our present study, we extended the use of base editors for direct correction of other β-thalassaemia mutations across multiple genomic loci such as the proximal promoter, exons, and introns of the *HBB* gene for the DSB free editing at the target site. We achieved highly efficient correction of disease-causing point mutations in the β-thalassaemic cellular model and confirmed the potential of base editing for therapeutic gene correction. The selection of base editors offers several advantages over conventional gene editing approaches. First, the base editors enable the generation of point mutations without introducing undesired insertions or deletions, which is especially advantageous in correcting mutations in the coding regions where the sequence integrity is critical. Secondly, highly efficient variants like ABE8e, which achieved more than 50% editing in almost all of the evaluated target sites. Further, the disadvantages associated with DNA double-strand break, such as deletions, inversion, translocations, p53 activation etc, could be reduced to a greater extent by the employment of base editors(15,16). This feature of base editors is crucial while targeting the mutations in homologous sequences of *HBB* and *HBD.* Besides these, base edited HSPCs have been demonstrated to engraft and maintain base conversion over time in *in-vivo* animal models to therapeutic potential (19,36).

We used an innovative strategy of introducing an engineered *HBB* construct harbouring diverse β-thalassemia mutations into the human erythroid cells for the primary validation of base editing components. This eliminated the requirement for patient samples or patient-derived cell models such as iPSCs with specific mutations to validate the gRNA and base editor. The editing frequency of gRNAs acquired using this approach was comparable to that achieved in the human β-thalassemia cellular model, indicating that this method is suitable and could be applied for other genetic diseases with a diverse mutation spectrum. Using the human erythroid cells containing the various β-thalassemia mutations, we validated gRNAs targeting seven functional (−88 C>T; −28A>G; initiation CD T>C; CD 15 G>A; IVSI-5 G>A; IVSI-110 G>A; IVS2-849 G>A) and one structural variant (CD26 G>A, HbE) β-thalassaemic/HbE mutations. Of the two ABE versions, ABE 8e consistently outperformed ABE7.10 in terms of on-target editing and expanded editing window, resulting in high bystander edits.

On the downside, nucleotide conversion using basic editors is confined to a small editing window, usually bases 3-9 (at the end distal to PAM). In addition, bystander editing of unintended nucleotides within the editing window is an unavoidable consequence. Careful designing of gRNA and the use of Cas9 variants recognising alternative PAM or with different editing windows could overcome these limitations to a great extent (40). The *HBB* −88(C>T) correction illustrates this point where only 20% on-target editing was obtained with ABE 7.10, while the use of the same gRNA with ABE 8e resulted in around 80% editing. Although we achieved substantial gene correction of two specific mutations in the *HBB* promoter and three *HBB* mutations in intronic regions, these edits co-occurred with bystander conversion, which were not reported to cause the disease phenotype. In addition, efficient base conversion at three exonic mutations was also accompanied by bystander edits within the editing window. Remarkably, most of the unintended modifications identified in the coding region of *HBB* were previously reported to display normal phenotypes in heterozygous conditions.

Furthermore, we demonstrated a simple and efficient approach for generating the human erythroid cellular model having β-thalassaemic point mutations at the *HBB* locus using base editors. We achieved intend ed editing at a significantly higher proportion than the bystander edits in almost all the sites in the generated human erythroid cellular model for β-thalassemia/HbE. However, it is critical to evaluate the functional β-haemoglobin synthesis, because all the variants formed due to bystander alterations may not be naturally occurring. In addition, it is essential to test the selected gRNA and base editor variants in patient-derived HSPCs to evaluate safety parameters such as off-target editing and multilineage engraftment potential before the gRNAs can be used for therapeutic application.

In conclusion, our study provides compelling evidence for using base editors to create and correct a spectrum of β-thalassemia mutations in human erythroid cells with high efficiency and precision, thus establishing a promising therapeutic framework to address the hereditary monogenic disorders caused by point mutations.

## Materials and method

### Designing of guide RNA, donor template and mutated *HBB* gene fragment

All guide RNAs (gRNAs) were designed using Snapgene software based on PAM availability and editing window. The gRNAs for electroporation were obtained from Synthego. For validation in cell lines, gRNAs were synthesised as oligonucleotides and cloned in the lentiviral backbone (#57822 or #57823). The ssODN donor template was designed manually with 181bps homology in 5’ and 3’ end (sequence mentioned in supplementary table 4) and was synthesised by Integrated DNA Technologies (IDT).

The mutations to be incorporated in the β-thalassemia gene fragment were chosen based on the prevalence, with particular attention to mutations with high frequency in India and worldwide.

The sequence of the *HBB* gene (Gene ID 3043) was downloaded from the NCBI website, and diverse β-thalassemia point mutations were shortlisted and incorporated into the sequence. The mutated gene fragment spanning 1800bp of the *HBB* gene, including the promoter and 3’UTR sequence, was obtained from IDT (sequence mentioned in supplementary table 5).

### Plasmid construct

Plasmids pMD2.G (Addgene #12259) and psPAX2 (Addgene# 12260) were a gift from Didier Trono) and were used for the virus production. pLKO5.sgRNA.EFS.GFP/RFP vectors (Addgene #57822 and #57823, respectively) were a gift from Benjamin Ebert (41) used for cloning guide RNAs. For preparing ABE, CBE, and ABE8e stables in *HBB* deleted HUDEP-2 cell harbouring β-thalassemia point mutations, lentivirus produced with pLenti-FNLS-P2A-Puro (Addgene#110841-CBE), pLenti-ABERA-P2A-Puro (Addgene#112675-ABE) gift from Lukas Dow (42) and pLenti-ABE8e-Puro were used. The pLenti-ABE8e-Puro plasmid was constructed from ABE8e (TadA-8e V106W) (Addgene#138495-ABE8e), a gift from David Liu (43), as previously reported (23).

### Cloning of gRNAs and *HBB* gene fragment harbouring β-thalassemia point mutations

The gRNAs were cloned into a lentiviral backbone with GFP/RFP marker as described previously (23) for validation in stable cell lines. Briefly, the gRNAs obtained as oligonucleotides were annealed and ligated into *BsmBI* (NEB) digested pLKO5.sgRNA.EFS.tRFP plasmid and transformed into DH10B competent cells and plated in LB-agar containing 100 μg/ml of Ampicillin. Obtained colonies were sequence confirmed to ensure that the guide RNAs sequence were inserted into the vector.

The gene fragment obtained from IDT was amplified using the primers listed in Table 2 (Mutated *HBB* gene F/R primers) and assembled into PpumI and XhoI digested pLKO5.sgRNA.EFS.GFP plasmid via Gibson Assembly using HiFi DNA assembly (NEB) as per the manufacturer’s protocol. The obtained clones were sequence confirmed via sanger sequencing.

### Cell culture

HEK293T cells were cultured in Dulbecco’s Modified Eagle Medium (DMEM, Gibco) supplemented with 10%FBS and 1% 1X Pen-Strep. HUDEP-2 cell line were cultured in Stemspan SFEM-II (StemCell Technologies) supplemented with 1 μM dexamethasone (Alfa Aesar), 1 μg/ml doxycycline (Sigma-Aldrich), 1X L-glutamine 200mM (Gibco™), 1X pen-Strep (Gibco™), SCF 50ng/ml (ImmunoTools) and 3 U/ml EPO (Zyrop 4000 IU injection) ( Kurita et al., 2013). HUDEP-2 cells were further differentiated into erythroid lineage using eight days differentiation protocol as previously described (23).

### Lentivirus Production

For virus production, 0.8 x10^6^ HEK 293T cells were seeded in one well of 6 well plates with DMEM (HyClone) supplemented with 10%FBS and 1% Pen-Strep. The transfection mixture was prepared by combining 0.5 μg pMD2.G, 0.5 μg psPAX2 (2^nd^ generation lentiviral packaging construct) and 1 μg (cloned gRNA or G-block) of a lentiviral vector with 6ul of FuGENE-HD in 100ul of OptiMem as per manufacturer’s instructions and added to the cells at 80% confluency. For preparing, *HBB* deleted HUDEP-2 stables virus was produced from two wells of a 6well plate for each construct (ABE7.10/ CBE / ABE8e). After 48hrs of transfection, the supernatant was collected and concentrated with Lenti-X Concentrator (Takara) and incubated at 4° C for 30 mins, then centrifuged at 4200 RPM for 30 minutes. The pellet was resuspended in 100μl of 1XPBS and stored at −80°C till use.

### Lentiviral Transduction

For gRNA validation, 0.25 x10^6 of *HBB* deleted HUDEP-2 cells in one well of 12 well plates were transduced with 100μl of the desired lentivirus along with 6μg/ml polybrene (Sigma-Aldrich) and 1% HEPES (1M buffer, Gibco). After 48hrs, the transduction efficiency was analysed by FACS by measuring GFP/RFP expression. For stable cell line preparation (ABE 7.10, CBE and ABE8e), the transduced cells were subjected to puromycin (1μg/ml (Sigma-Aldrich)) selection for 10 days.

### Electroporation in HUDEP-2 cell line

For the creation of the HbE disease model, HDR was performed by electroporation of RNP complex (Cas9: 50pmole; sgRNA: 100pmole) along with donor template(100pmole) in 1x 10^6 HUDEP-2 cells using Lonza 4d electroporator (Pulse code CA137). For the generation of the Initiation Codon and IVS2-849 disease model 1x 10^6, HUDEP-2 cells were electroporated with respective sgRNA (100pmole) and ABE mRNA (5μg) using Lonza 4d electroporator with (Pulse code EN138). 96 hrs post-electroporation editing was evaluated by sanger sequencing.

### Single-cell sorting

To obtain a homozygous and heterozygous clone for Initiation Codon (T->C), IVS2-849(A->G) and HbE (G->A) mutation, edited HUDEP-2 cells were sorted in 96well plate using FACS (BD - FACSAria™ III Cell Sorter). The sorted cells were expanded further, and edited alleles were verified by sanger sequencing.

### Analysis of editing efficiency

Genomic DNA was isolated using DNA isolation Kit (Nucleospin Blood-Macherey-Nagel) and Sanger sequencing was performed by amplifying the target regions with GoTaq^®^ Hot Start Polymerase Premix (Promega) with the respective primers as mentioned in Table 2. Sanger sequencing data were analysed for base editing efficiency by EditR and indels by Inference of CRISPR Edits (ICE) (Synthego) (44,45).

### Evaluation of *HBB* gene by Quantitative Real-Time PCR Analysis

To quantify the *HBB* gene deletion in the HUDEP-2 cell line, a quantitative real time-PCR was carried out as previously reported (46) performed by using genomic DNA with SYBR Premix Ex Taq II (Takara Bio Inc.) as per the manufacturer’s protocol using specific primers as mentioned in Table 2.

### Evaluation of *HBB* gene deletion by PCR

To quantify the *HBB* gene deletion in HUDEP-2 cells, genomic DNA was isolated from *HBB* null cells using DNA isolation Kit (Nucleospin Blood-Macherey-Nagel) and PCR was performed using primers (Table 2) with PrimeSTAR^®^ GXL DNA Polymerase (Takara).

### MLPA

Genomic DNA was isolated from HUDEP-2 cells, HUDEP-2 *HBB* null cells and HUDEP-2 with mutated *HBB* gene cells using DNA isolation Kit (Nucleospin Blood-Macherey-Nagel) and MLPA (Multiplex Ligation-dependent Probe Amplification) was performed using SALSA MLPA probe mix P102 *HBB* with 100ng/μl DNA as per the manufacturer’s protocol.

### mRNA synthesis

ABE 8e mRNA was prepared using T7 mScript™ Standard mRNA Production System (CELLSCRIPT−) as per the manufacturer’s protocol.The template for IVT was prepared by digesting ABE8e (TadA-8e V106W) plasmid (Addgene# 138495-ABE8e) using Pme1 restriction endonuclease (NEB).

### Statistics

GraphPad Prism 8.1 has been used for the statistical tests.

## Acknowledgements

The research reported in this work was supported by NAHD grant: BT/PR17316/MED/31/326/2015 (Department of Biotechnology, New Delhi, India), EMR grant: EMR/2017/004363 (Science and Engineering Research Board [SERB], New Delhi, India), Indo-US GET in Fellowship_2018_066 (Indo-US Science & Technology Forum [IUSSTF]), and DBT grant: BT/PR38392/GET/119/301/2020. We sincerely acknowledge CSCR (a unit of inStem, CMC Campus, Vellore, India) for providing startup funds. Anila George and Nithin Sam Ravi are supported by Senior Research Fellowship from the Council of Scientific & Industrial Research India. Vignesh Rajendiran is supported by Senior Research Fellowship DBT India. We thank Mr Dhiyaneshwaran.S for helping us in making figures and Dr. Muthuraman. N. for proofreading the manuscript. We also acknowledge CSCR Core for all instrumentation facilities and support while performing experiments.

## Authorship Contribution

MKM conceived and supervised the study; KP performed most of the experiment under the guidance of MKM; KP and MKM analysed the data; VR, AG, NSR, DN and AAP provided experimental support; KL and DN assisted in screening most prevalent β-thalassaemic mutations; GM and YP under the guidance of SM assisted in ABE8e mRNA preparation; VV under the guidance of ST assisted in performing HDR; YN and RK provided HUDEP-2 cells; KP, LP, DN, NSR and MKM prepared all figures; KP, AG, DN and MKM wrote the manuscript; AS, RVS, ST, SM and BP provided intellectual insight and feedback on the data and manuscript. All authors contributed to manuscript revision, read, and approved the submitted version.

## Notes

### Competing Interest Statement

The authors have declared no competing interest.

