## Supplementary file for "Precise modelling and correction of a spectrum of β-thalassemic mutations in human erythroid cells by base editors"

Supplementary figure 1

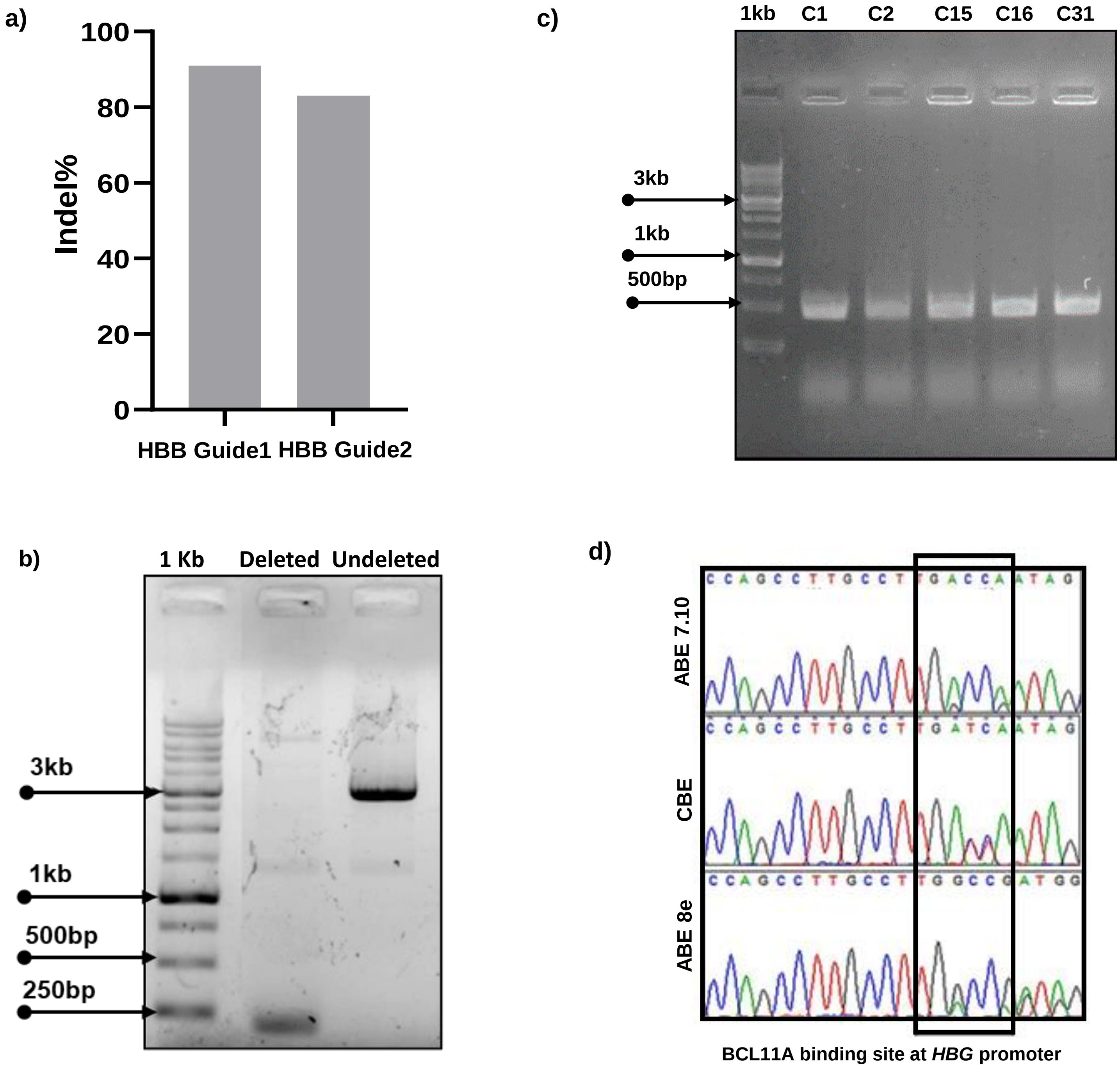

**Supplementary figure 1: Characterisation of *HBB* null HUDEP-2 cells expressing base editor variants: a)** Editing efficiency of gRNAs used for *HBB* deletion in HUDEP-2 cells determined by Sanger sequencing. **b)** *HBB* deletion reconfirmed using a different set of oligos in *HBB* null HUDEP-2 cells; the amplified band at 232bp and 3330bp represents the deleted and control samples, respectively. **c)** Confirmation of homozygous *HBB* deleted single-cell clones using PCR (band at 500bp represents the deleted samples). **d)** Validation of editing efficiency in engineered HUDEP-2 cells expressing base editor variants.

### Supplementary figure 2

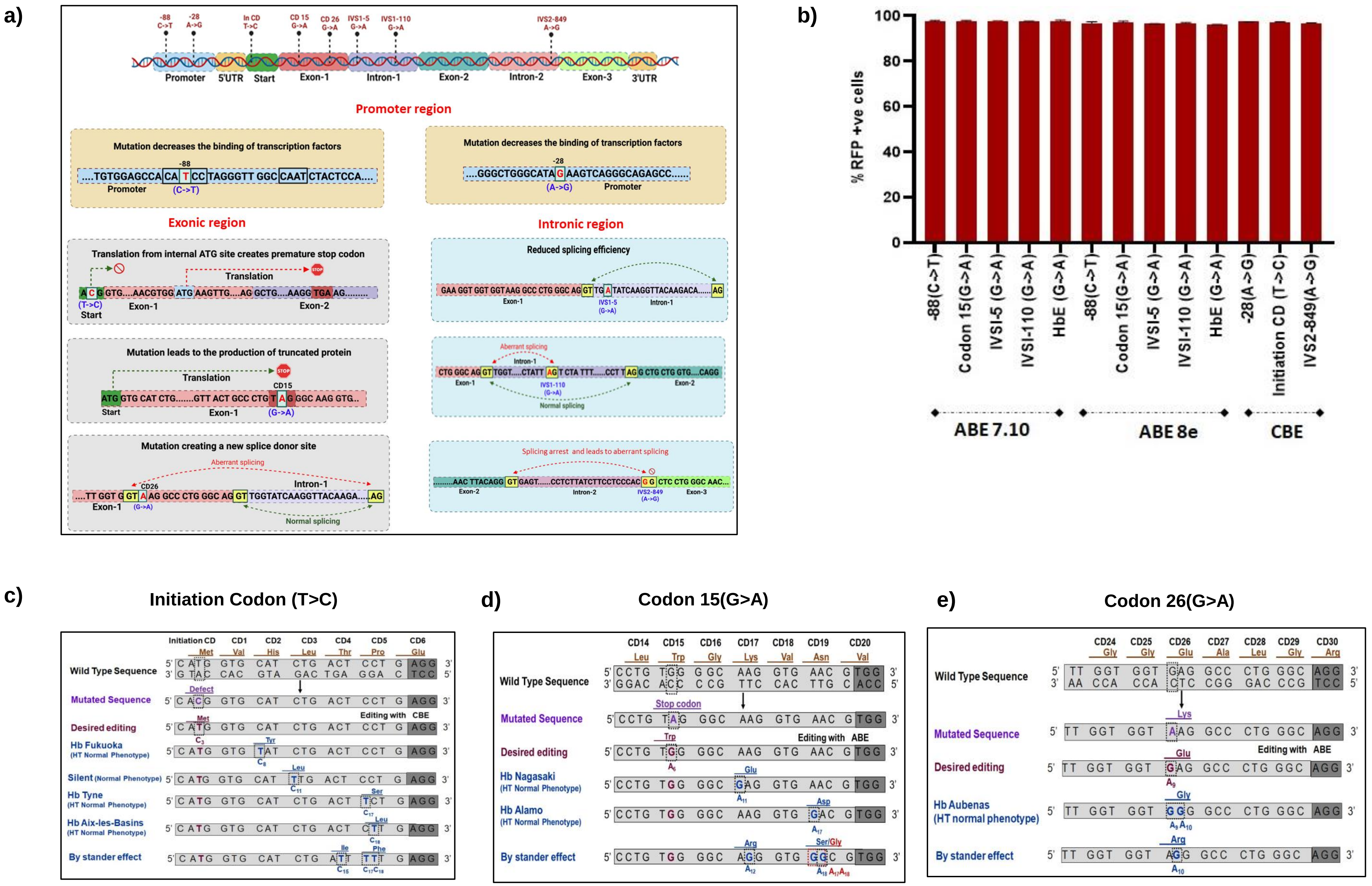

**Supplementary figure 2: Mechanism of β-thalassemia point mutations and their correction using respective gRNAs in engineered HUDEP-2 cells using base editors a)** A brief mechanism of the eight different β-thalassemia point mutations that are corrected using base editors in this study are graphically represented. **b)** Transduction efficiency of gRNAs targeting various *HBB* mutations in engineered HUDEP-2 cells. **c-e)** Pictorial representation of possible *HBB* variants that could arise due to the bystander edits while targeting the HBB exonic region.

**Supplementary table 1: gRNAs for HBB gene deletion**

| S.No. | Guide RNA | PAM | Sequence (5' -> 3’) |
|---|---|---|---|
| 1. | *HBB* Guide 1 | NGG | ACTATCAATGGGGTAATCAG |
| 2. | *HBB* Guide 2 | NGG | ATCTCGCCGTAAAACATGGA |

**Supplementary table 2: List of primers**

| S.No. | Primer Name | Tm | Amplicon size | Sequence (5' -> 3’) |
|---|---|---|---|---|
| **I** | ***HBB* gene deletion** | | | |
| 1. | *HBB* gene deletion primer (F) | 60 | 3300 /500 bps | CGATCACGTTGGGAAGCTAT |
| | *HBB* gene deletion primer (R) | | | CCAGTCCTTCCAAAGCAGAC |
| 2. | *HBB* gene G1 (F) | 59 | 638 bps | GATGGGAGAAAGGCGATCAC |
| | *HBB* gene G1 (R) | | | AAACTCTACCTCGGTTCTAAGC |
| 3. | *HBB* gene G2 (F) | 59 | 460 bps | CTAATGCACATTGGCAACAGC |
| | *HBB* gene G2 (R) | | | TTTCATACTAAGCCCAGTCCTTC |
| 4. | *HBB* deletion PCR (F) | 60 | 232 bps | CTTTTCCCCTCCTACCCCTA |
| | *HBB* deletion PCR (R) | | | TCAAACCATGACCCCTGTTT |
| 5. | RT -PCR undeleted (F) | - | 234 bps | ACTCCTAAGCCAGTGCCAGA |
| | RT -PCR undeleted (R) | | | CTCAGGAGTCAGATGCACCA |
| **II** | **Correction of the spectrum of β-thalassemia mutations integrated in HUDEP-2 HBB null cells:** | | | |
| 1. | Mutated *HBB* gene (F) | 57 | 1880 bps | CGTTTCAGACCCACCTCCCA |
| | Mutated *HBB* gene (R) | | | GCTCTGCCCACTGACGGGCA |
| 2. | Init CD - IVS1-110 (F) | 57 | 857 bps | GTGAATAGAGTTAGGCAGGG |
| | Init CD - IVS1-110 (R) | | | CTGAGACTTCCACACTGATGC |
| 3. | IVS2-849 (F) | 57 | 833 bps | CACATATTGACCAAATCAGGG |
| | IVS2-849 (R) | | | CGACATCACTTTCCCAGT |
| **III** | **Creation & Correction of β-thalassemia mutations:** | | | |
| 1. | Initiation Codon / HbE (F) | 55 | 706 bps | GTCATCACTTAGACCTCACC |
| | Initiation Codon/ HbE (R) | | | CTGTACCCTGTTACTTATCC |
| 2. | IVS2-849 (F) | 56 | 1153 bps | CACATATTGACCAAATCAGGG |
| | IVS2-849 (R) | | | TCAAACCATGACCCCTGTTT |

**Supplementary table 3: List of guide RNAs**

| S.No. | Guide RNA | PAM | Sequence (5' -> 3’) |
|---|---|---|---|
| **I** | **Correction of a spectrum of β-thalassemia mutations integrated in HUDEP-2 *HBB* null cells:** | | |
| 1. | -88 (C->T) | AGG | GGATGTGGCTCCACAGGGTG |
| 2. | -28(A->G) | TGG | CTGACTTCTATGCCCAGCCC |

| | | | |
|---|---|---|---|
| 3. | Initiation Codon(T->C) | AGG | CACGGTGCATCTGACTCCTG |
| 4. | CD15 (G->A) | TGG | CCTGTAGGGCAAGGTGAACG |
| 5. | HbE gRNA1(G->A) | AGG | TGGTAAGGCCCTGGGCAGGT |
| 6. | HbE gRNA2(G->A) | AGG | TTGGTGGTAAGGCCCTGGGC |
| 7. | IVS1-5(G->A) | AGG | TGATATCAAGGTTACAAGAC |
| 8. | IVS1-110(G->A) | AGG | TAGTCTATTTTCCCACCCTT |
| 9. | IVS2-849 (A->G) | AGG | AGCCGTGGGAGGAAGATAAG |
| **II** | **Creation of β-thalassemia mutations in HUDEP-2 cells:** | | |
| 1. | Initiation Codon (ATG->ACG) | AGG | TGCACCATGGTGTCTGTTTG |
| 2. | IVS2-849 (A->G) | TGG | ACAGCTCCTGGGCAACGTGC |
| 3. | HbE HT gRNA1 | AGG | AAGGTTACAAGACAGGTTTA |
| 4. | HbE HO gRNA2 | AGG | CGTGGATGAAGTTGGTGGTG |

**Supplementary table 4: Sequence of donor template**

| **S.No.** | **Name** | **Sequence (5'->3')** |
|---|---|---|
| 1. | HbE_ssODN (181bps) | Caaacagacaccatggtgcatctgactcctgaggagaagtctgccgttactgccctgtggggcaaggtg aacgtggatgaagttggtggtAaggccctgggcagGTTGGTATCAAGGTTACAAGACAGGT TTAAAGAGACCAATAGAAACTGGGCATGTGGAGACAGAGAAGACTCTTGGG |

**Supplementary table 5: Mutated HBB sequence**

| **S.No.** | **Name** | **Sequence (5'->3')** |
|---|---|---|
| 1. | Mutated HBB (1880pbs) | cgtttcagacccacctcccaaccccgaggggacccGCCAGTGCCAGAAGAGCCAAGGACAG GTACGGCTGTCATCACTTAGACCTCACCCTGTGGAGCCACATCCTAGGGTTGGC CATTCTACTCCCAGGAGCAGGGAGGGCAGGAGCCAGGGCTGGGCATAGAAGT CAGGGCAGAGCCATCTATTGCTtCcatttgcttctgacacaactgtgttcactagcaacctcaaa cagacaccaCggtgcatctgactcctgaggagaagtctgccgttactgccctgtAgggcaaggtgaacg tggatTaagttggtggtAaggccctgggcagGTTGATATCAAGGTTACAAGACAGGTTTAA GGAGACCAATAGAAACTGGGCATGTGGAGACAGAGAAGACTCTTGGGTTTCTG ATAGGCACTGACTCTCTCTGCCTattAgtctattttcccacccttAggctgctggtggtctaccct tgAacccagaggttctttgagtcctttggggatctgtccactcctgatgctgttatgggcaaccctTaggtg aaggctcatggcaagaaagtgctcggtgcctttagtgatggcctggctcacctggacaacctcaagggca cctttgccacactgagtTagctgcactgtgacaagctgcacgtggatcctgagaacttcagggtgaCtct atgggacgcttgatgttttctttccccttcttttctatggttaagttcatgtcataggaaggggataagtaac agggtacagtttagaatgggaaacagacgaatgattgcatcagtgtggaagtctcaggatcgttttagttt cttttatttgctgttcataacaattgttttcttttgtttaattcttgctttcttttttttcttctccgcaatttttac tattatacttaatgccttaacattgtgtataacaaaaggaaatatctctgagatacattaagtaacttaaa aaaaaactttacACAGTCTGCCTAGTACATTACTATTTGGAATATATGTGTGCTTATT TGCATATTCATAATCTCCCTACTTTATTTTCTTTTATTTTTAATTGATACATAATCA TTATACATATTTATGGGTTAAAGTGTAATGTTTTAATATGTGTACACATATTGAC |

| | | |
|---|---|---|
| | | CAAATCAGGGTAATTTTGCATTTGTAATTTTAAAAAATGCTTTCTTCTTTTAATATACTTTTTTGTTTATCTTATTTCTAATACTTTCCCTAATCTCTTTCTTTCAGGGCAATAATGATACAATGTATCATGCCTCTTTGTACCATTCTAAAGAATAACAGTGATAATTTCTGGGTTAAGGTAATAGCAATATCTCTGCATATAAATATTTCTGCATATAAATTGTAACTGAGGTAAGAGGTTTCATATTGCTAATAGCAGCTACAATCCAGGTACCATTCTGCTTTTATTTTATGGTTGGGATAAGGCTGGATTATTCTGAGTCCAAGCTAGGCCCTTTTGCTAATCATGTTCATACCTCTTATCTTCCTCCCACGgctcctgggcaacgtgctggCctgtgtgctggcccatcactttggcaaaTaattcaccccaccagtgcaggctgcctatcagaaagtggtggctggtgtggctaatgccctggcccacaagtatcactaagctcgctttcttgctgtccaatttctattaaaggttcctttgttccctaagtccaactactaaactgggggatattatgaagggccttgagcatctggattctgcctaatGaaaaacatttattttcattgcaaTGATGTATTTAAATTATTTCTGAATATTTTACTAAAAAGGGAATGTGctcgagtggctccggtgcccgtcagtgggcagagc |
